# Altered Bile Acid Profile Associates with Cognitive Impairment in Alzheimer’s Disease – An Emerging Role for Gut Microbiome

**DOI:** 10.1101/281956

**Authors:** Siamak MahmoudianDehkordi, Matthias Arnold, Kwangsik Nho, Shahzad Ahmad, Wei Jia, Guoxiang Xie, Gregory Louie, Alexandra Kueider-Paisley, M. Arthur Moseley, J. Will Thompson, Lisa St John Williams, Jessica D. Tenenbaum, Colette Blach, Rebecca Baillie, Xianlin Han, Sudeepa Bhattacharyya, Jon B. Toledo, Simon Schafferer, Sebastian Klein, Therese Koal, Shannon L. Risacher, Mitchel Allan Kling, Alison Motsinger-Reif, Daniel M. Rotroff, John Jack, Thomas Hankemeier, David A. Bennett, Philip L. De Jager, John Q. Trojanowski, Leslie M. Shaw, Michael W. Weiner, P. Murali Doraiswamy, Cornelia M. van Duijn, Andrew J. Saykin, Gabi Kastenmüller, Rima Kaddurah-Daouk, for the Alzheimer’s Disease Neuroimaging Initiative and the Alzheimer Disease Metabolomics Consortium

## Abstract

**Introduction:** Increasing evidence suggests a role for the gut microbiome in central nervous system disorders and specific role for the gut-brain axis in neurodegeneration. Bile acids (BA), products of cholesterol metabolism and clearance, are produced in the liver and are further metabolized by gut bacteria. They have major regulatory and signaling functions and seem dysregulated in Alzheimer disease (AD).

**Methods:** Serum levels of 15 primary and secondary BAs and their conjugated forms were measured in 1,464 subjects including 370 cognitively normal older adults (CN), 284 with early mild cognitive impairment (MCI), 505 with late MCI, and 305 AD cases enrolled in the AD Neuroimaging Initiative. We assessed associations of BA profiles including selected ratios with diagnosis, cognition, and AD-related genetic variants, adjusting for cofounders and multiple testing.

**Results:** In AD compared to CN, we observed significantly lower serum concentrations of a primary BA (cholic acid CA) and increased levels of the bacterially produced, secondary BA, deoxycholic acid (DCA), and its glycine and taurine conjugated forms. An increased ratio of DCA:CA, which reflects 7α-dehydroxylation of CA by gut bacteria, strongly associated with cognitive decline, a finding replicated in serum and brain samples in the Rush Religious Orders and Memory and Aging Project. Several genetic variants in immune response related genes implicated in AD showed associations with BA profiles.

**Conclusion:** We report for the first time an association between altered BA profile, genetic variants implicated in AD and cognitive changes in disease using a large multicenter study. These findings warrant further investigation of gut dysbiosis and possible role of gut liver brain axis in the pathogenesis of AD.

## 1. Introduction

Alzheimer’s disease (AD), a progressive neurodegenerative disorder, is the leading cause of dementia in old age affecting over 40 million people worldwide[1]. There are currently no therapies to prevent or slow AD progression, highlighting our incomplete knowledge of disease mechanisms and the need for new drug targets. A large number of biochemical processes are affected in AD and genes implicated in AD highlight the possible roles for lipid processing, immune function, phagocytosis, (innate) immunity and neurotransmitter function, biological pathways that may affect metabolism[2, 3]. Recent AD hypotheses implicate viral and bacterial contributions to disease pathogenesis[4-6].

Bidirectional biochemical communication between the brain and the gut contribute to a variety of neurodegenerative and psychiatric diseases[7-10]. The gut microbiome and the host collaboratively produce a large array of small molecules that impacts human health[11, 12]. Recently, a role for the gut microbiome in motor dysfunction in Parkinson’s disease has been highlighted[13] and several animal models of AD showed a possible role of gut bacteria in amyloid-beta (Aβ) pathology[14, 15]. The APP transgenic mouse model of AD had a drastically altered gut microbiome composition compared to wild-type mice[15]. Other studies linked pro-inflammatory bacteria, such as gram-negative producers of neurotoxic lipopolysaccharides, to brain amyloidosis and systemic inflammation, a central feature of AD[16, 17]. These studies suggest microbial dysbiosis or imbalance could potentially contribute to AD pathogenesis.

Cholesterol metabolism in the liver is thought to play a key role in AD[18]. In fact, many cholesterol metabolism genes (e.g., *BIN1, CLU, PICALM, ABCA7, ABCG1*, and *SORL1)* are among the top AD susceptibility loci identified by genome-wide association studies[2, 19]. Cholesterol is cleared through production of bile acids (BAs). Primary BAs, chenodeoxycholic acid (CDCA) and cholic acid (CA), are synthesized from cholesterol in the liver, conjugated with glycine or taurine, secreted into the gallbladder via the bile salt export pump (BSEP), and transported to the intestine to be metabolized by gut bacteria (**Fig. 1**). Intestinal anaerobic bacteria deconjugate the liver-derived BAs through the action of bile salt hydrolases (BSH) to their respective free BAs. Subsequently, anaerobe bacteria convert primary BAs to the secondary BAs. That is, CA is converted to deoxycholic acid (DCA). CDCA is converted to lithocholic acid (LCA) and ursodeoxycholic (UDCA) through 7α or 7β –dehydroxylation, respectively[20, 21]. In the terminal ileum and colon, BAs are reabsorbed by the enterocytes and released into the portal vein for return to the liver where they are conjugated to produce their glycine and taurine forms.

**Fig. 1A.**
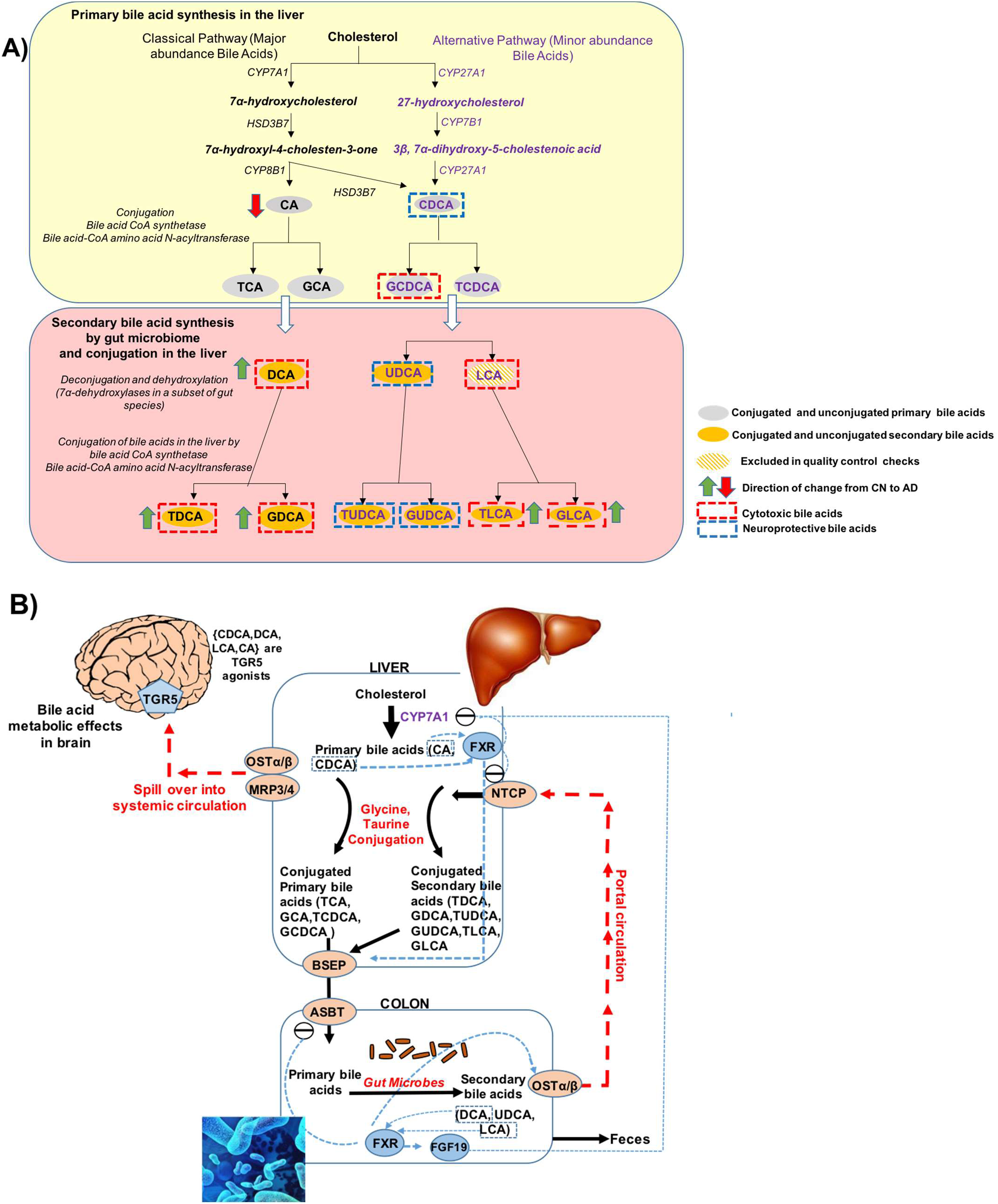
Bile acid synthesis and cholesterol clearance pathway. B. Regulation of bile acid synthesis by feedback mechanism and bile acid transport through enterohepatic circulation. In the liver the bile acids (CDCA, DCA, LCA, CA) activate FXR that inhibits (via a repressor SHP, not shown here) the rate-limiting enzyme CYP7A1. The bile acids via FXR/SHP also inhibit the influx transporter NTCP; induce BSEP and canalicular bile acid secretion. In the intestine, bile acids, via FXR, inhibit the uptake transporter ASBT, decreasing absorption and increasing basolateral secretion into portal circulation by inducing OSTα & β. Bile acid activated FXR in the intestine also exerts inhibition on CYP7A1 in the liver via FGF19 pathway. At the basolateral membrane of hepatocytes, transporters OSTα & β, and also MRP3 and MRP4, secrete bile acids into the systemic circulation. *Abbreviations:* ASBT: Apical Sodium-dependent Bile acid Transporters; BSEP: Bile Salt Export Pump; FXR: Farnesoid X Receptor; NTCP: Sodium/Taurocholate Co-transporting Polypeptide; SHP: Small heterodimer partner.

Beyond BAs role in cholesterol clearance, BAs are major regulators for maintaining energy homeostasis through binding to nuclear receptors, including FXR, among others. BAs also modulate the gut microbiome[22, 23]. Both primary and secondary BAs are present in the brains of mice and humans with evidence that they cross the blood-brain barrier[24-29]. Some BAs such UDCA exert beneficial effects while others are known to be cytotoxic[30-34]. In particular, DCA’s toxicity has been associated with modulating apoptosis involving mitochondrial pathways in a variety of tissues and cell types[35-37].

In recent pilot human studies, BA profiles were shown to be affected in AD[26, 38-40]. Here, we used a targeted metabolomics approach to evaluate BA profiles in a large cohort of 1,464 individuals enrolled in the AD Neuroimaging Initiative (ADNI) where rich clinical, imaging, and genetic data exist. We used this data to address the following:

1. Investigate if BA profiles are altered in MCI and AD patients and if these differences are related to cognitive decline.
2. Use ratios of BAs to pinpoint possible enzymatic alterations in the liver and in the gut microbiome that directly contribute to altered BA profile.
3. Investigate whether immune related AD genome-wide significant genes affect levels of BAs in circulation as markers for altered gut microbiome function.

In a subsequent paper we evaluated correlations between BAs and ATN biomarkers of AD including CSF biomarkers, atrophy, and brain glucose metabolism.

## 2. Methods

### 2.1. Study cohorts and samples

#### 2.1.1. ADNI baseline samples

Data used in the preparation of this article were downloaded from the ADNI database (http://adni.loni.usc.edu/). The ADNI studies have recruited over 1,500 adults, ages 55 to 90, consisting of cognitively normal older individuals (CN), individuals with subjective memory concerns (SMC), subjects with early (EMCI) or late mild cognitive impairment (LMCI), and patients with early probable AD dementia. Subjects categorized as SMC were excluded in this study. For key clinical and demographic variables of ADNI participants included in this study, see **Table 1.**

**Table 1.A.** Demographics of ADNI participants stratified by baseline diagnosis^a^.

| Variable | N | CN<br>(N=370) | EMCI<br>(N=284) | LMCI<br>(N=505) | AD<br>(N=305) | *p*-value<sup>b</sup> |
| --- | --- | --- | --- | --- | --- | --- |
| Age | 1464 | 74.58(5.71) | 71.12(7.51) | 73.95(7.59) | 74.70(7.79) | ***P*<0.001** |
| Sex: Female, No. (%) | 1464 | 190(51%) | 130(46%) | 197(39%) | 139(46%) | ***P*=0.004** |
| Education, years | 1464 | 16.28(2.92) | 15.95(2.66) | 15.87(2.90) | 15.16(3.00) | ***P*<0.001** |
| BMI (Kg/M<sup>2</sup>) | 1461 | 27.05(4.46) | 28.06(5.41) | 26.54(4.25) | 25.83(4.69) | ***P*<0.001** |
| ≥1 *APOE* ε4 allele, No. (%) | 1464 | 104(28%) | 121(43%) | 273(54%) | 202(66%) | ***P*<0.001** |
| ADAS-Cog13<sup>c</sup> | 1455 | 9.19(4.17) | 12.64(5.40) | 18.67(6.62) | 29.67(8.20) | ***P*<0.001** |

<sup>a</sup>Data are reported as mean(SD) unless otherwise indicated. Bolded values indicate statistical significance. SD: Standard deviation.
<sup>b</sup>Based on 2-sample *t* tests, or Pearson $\chi^2$ tests.
<sup>c</sup>Score explanations: ADAS-Cog13 range, 0 (best) to 85 (worst).
*Abbreviations:* AD: Alzheimer's disease; BMI: Body mass index; CN: Cognitively normal; EMCI: Early mild cognitive impairment; LMCI: Late mild cognitive impairment; ADAS-Cog13, Alzheimer Disease Assessment Scale 13-item cognitive subscale.

**Table 1.B.** Demographics of ADNI participants stratified by MCI progression to AD^a^.

| Variable | N | MCI-NonConverter<br>(N=538) | MCI-Converter<br>(N=251) | *p*-value<sup>b</sup> |
| --- | --- | --- | --- | --- |
| Age | 789 | 72.47(7.90) | 73.91(7.08) | ***P*=0.01** |
| Sex: Female, No. (%) | 789 | 41% (223) | 41% (104) | ***P*=1** |
| Education, years | 789 | 15.95(2.85) | 15.79(2.76) | ***P*=0.43** |
| BMI (Kg/M<sup>2</sup>) | 788 | 27.37(4.80) | 26.47(4.61) | ***P*=0.005** |
| ≥1 *APOE* ε4 allele, No. (%) | 789 | 41%(223) | 68%(171) | ***P*<0.001** |
| ADAS-Cog13 | 786 | 14.26(6.04) | 21.31(5.94) | ***P*=0.29** |

<sup>a</sup>MCI Subjects that converted to AD dementia in 4 years after baseline were labeled as MCI Converter.

#### 2.1.2. The Religious Orders Study and the Rush Memory and Aging Project (ROS/MAP) for replication of key finding

The ROS/MAP studies are both longitudinal cohort studies of aging and AD at Rush University, and are designed to be used in joint analyses to maximize sample size. ROS enrolled individuals from religious orders (nuns, priests, brothers) across the United States[41]. MAP was designed to complement the ROS study by using a similar structure and design as ROS, but enrolling participants with a wider range of life experiences and socioeconomic status from the Chicago, IL metropolitan area[42]. The entire ROS/MAP cohort consists of approximately 3,300 participants, more than 1,500 of whom have come to autopsy (www.radc.rush.edu). We measured a subset of serum BAs in 566 subjects (446 CN, 109 MCI, and 11 AD), as well as a subset of BAs in postmortem brain samples of 111 subjects with brain pathology performed (51 CN, 31 MCI, and 27 AD at time of death), of whom 93 also had serum measurements. Key demographic variables of ROS/MAP cohort are listed in **Supplementary Table 1**.

#### 2.1.3. Rotterdam study (RS)

RS was used to perform association of BAs with AD genetic variants. RS is a prospective population based study[43]. At the baseline examination in 1990-93, 7983 subjects ≥ 55 years of age were recruited from the Ommoord district of Rotterdam (RS-I). All the study participants were extensively interviewed and physically examined at baseline and after every 3 to 4 years. During 2000 to 2001, the baseline cohort (RS-I) was expanded with 3011 subjects ≥55 years of age, who were not yet part of RS-I (RS-II). In this analysis, fasting serum BAs were measured for 488 dementia-free subjects with mean(SD) age of 73.1(6.3) from RS-I using Metabolon platform (Durham, North Carolina, USA) as described previously[44] (see **Supplementary Table 2** for demographics).

### 2.2. Sample collection and quantification of BAs

Targeted metabolomics profiling was performed to measure concentrations of 20 BA metabolites in serum samples of the ADNI cohorts (see **Table 2** for list of BAs and abbreviations). Morning fasting serum samples from the baseline visit were collected and aliquoted as described in the ADNI standard operating procedures. BA quantification was performed by liquid chromatography tandem mass spectrometry using the Biocrates^®^ Life Sciences Bile Acids Kit (BIOCRATES Life Science AG, Innsbruck, Austria) according to manufacturer’s instructions.

**Table 2.**
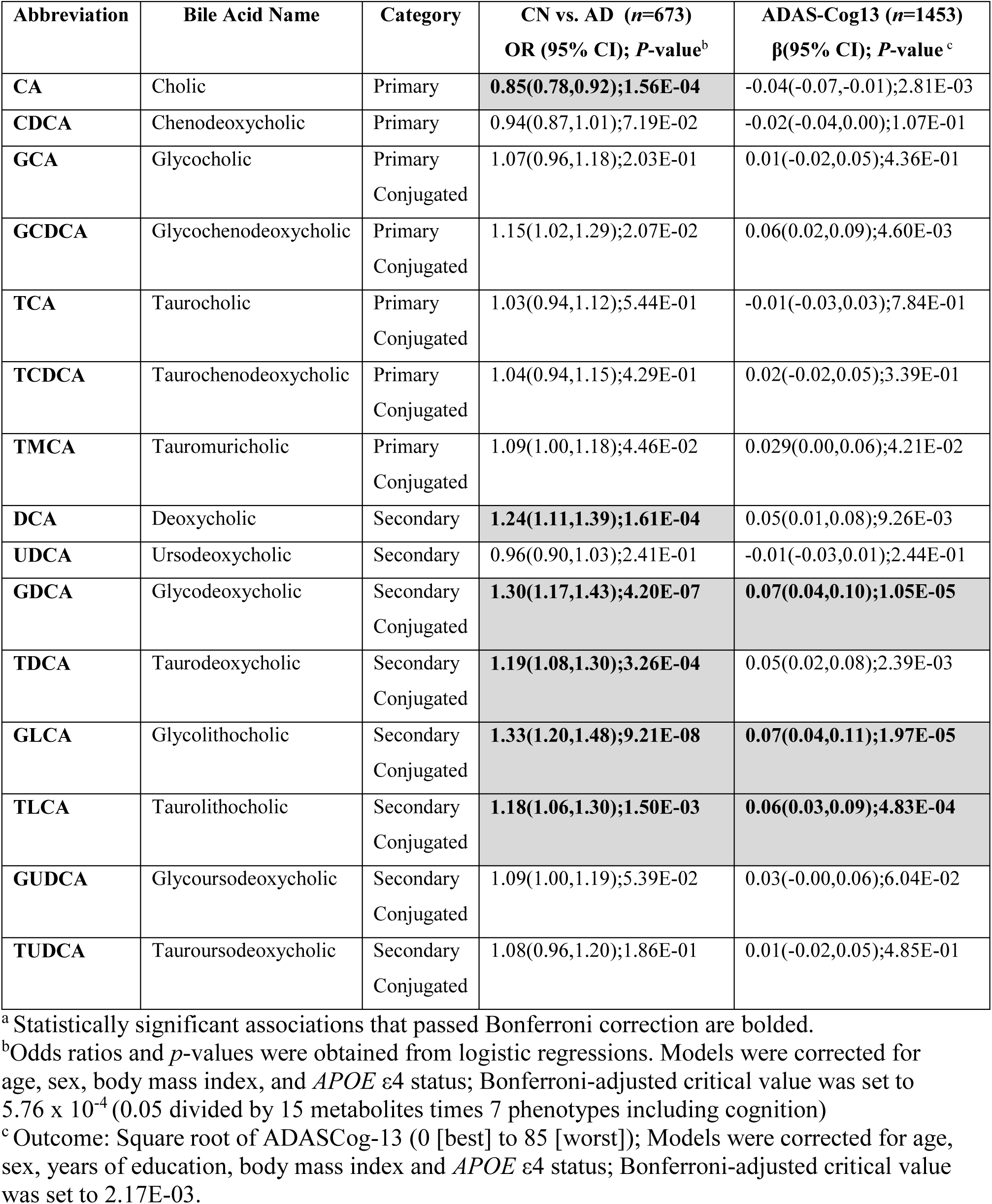
Primary and Secondary bile acids measured in the ADNI study and their cross-sectional association with diagnosis and cognition^a^.

In the ROS/MAP, quantification of BA concentration in 566 serum samples and 111 postmortem brain samples was performed at the University of Hawaii cancer center using ultra-performance liquid chromatography coupled to a tandem mass spectrometry (UPLC-MS/MS) system (ACQUITY UPLC-Xevo TQ-S, Waters Corp., Milford, MA)[45].

In the RS study, serum BAs were measured in 488 serum samples using the non-targeted Metabolon platform (Durham, North Carolina, USA).

### 2.3. Quality control of BA profiles

Metabolomics lab staff were blinded to diagnosis and pathological data in all the studies. In ADNI, after unblinding and data release, metabolite profiles went through quality-control (QC) checks and data preprocessing including batch-effect adjustment, missing value imputation, and log-transformation (**Supplementary Methods** and **Supplementary Table 3).** After QC correction, the dataset included 15 BAs (5 BAs did not pass QC criteria) for a total of 1,464 subjects (after excluding 99 SMC). The preprocessed BA values after QC were used for subsequent association analyses directly or were adjusted to take into account the effect of medications on BA levels[46]. The list of medications selected for adjustment for each BA is shown in **Supplementary Table 7**. We performed all analyses using both medication adjusted and unadjusted BA levels, results derived from medication-adjusted data and the adjustment process are described in **Supplementary Methods** and its accompanying tables.

In both RS and ROS/MAP, missing metabolite levels were imputed using half of the limit of detection. Log-transformed values were used in subsequent analyses.

### 2.4. Clinical Outcomes

For ADNI data, continuous response variables included the modified Alzheimer Disease Assessment Scale 13-item cognitive subscale (ADAS-Cog13; range, 0 [best] to 85 [worst] points), an index of general cognitive functioning. Categorical response variables included clinical diagnosis at baseline and MCI conversion (MCI-NonConverter, MCI-Converter). For the ROS/MAP cohort, cognition was measured using a battery of tests (details are published[47-50]). A composite measure of global cognition was created by averaging the z-scores of all tests as previously described[50]. Mean and standard deviation at baseline were used to compute z-scores. A negative z-score means that an individual has an overall score that is lower than the average of the entire sample at baseline. Cognitive tests were used from the same cycle as serum, and proximate to death for brain.

### 2.5. Genotype and whole genome sequencing data

#### Whole genome sequencing

For 817 ADNI participants, whole-genome sequencing (WGS) was performed on blood-derived genomic DNA. Samples were sequenced on the Illumina HiSeq2000 using paired-end read chemistry and read-length of 100bp at 30–40X coverage. For data processing and QC, an established analysis pipeline based on GATK was used. The QC steps included participant sex check, participant identity check, and variant quality check of the Illumina-generated VCF files (see Saykin et al., 2015 for details[51]).

#### DNA genotyping

in the participants of the RS cohort was performed using 550K, 550K duo, or 610K Illumina arrays at the internal genotyping facility of Erasmus Medical Center, Rotterdam. Study samples with excess autosomal heterozygosity, call rate < 97.5%, ethnic outliers and duplicate or family relationships were excluded during quality control analysis. Genotype exclusion criteria further included call rate < 95%, Hardy-Weinberg equilibrium *p* < 1.0×10^−6^ and Minor Allele Frequency (MAF) < 1%. Genetic variants were imputed to the Haplotype Reference Consortium (HRC) reference panel (version 1.0)[52] using the Michigan imputation server[53].

#### Reference genetic associations

with BA profiles in healthy individuals were obtained from supplementary data of the atlas of genetic influences on blood metabolites[44]. To obtain genome-wide genetic associations with DCA, we considered all suggestive significant results with *P* < 1.0 × 10^−5^. Gene and complex trait annotations of the 13 resulting genetic loci were performed using the SNiPA tool v3.2[54] and the NHGRI-EBI Catalog of published genome-wide association studies (www.ebi.ac.uk/gwas; accessed 02/01/2018, version 1.0)[55]. Lookup of AD genetic associations for DCA candidate variants was performed using the IGAP repository[2].

### 2.6. Statistical analysis

Differences of demographic, clinical, and cognitive measurements among the clinical diagnostic groups were evaluated using 2-sample *t*-test (for continuous variables) and Pearson Chi-squared test (for categorical variables). All analyses were performed in a metabolite-wise manner and Bonferroni-adjusted critical values were used to assess statistical significance. All models included age at baseline, sex, *APOE* ε4, and log_10_-transformed body mass index (BMI). For cognition, number of years of education was added as an additional covariate.

Separate binary logistic regression models were conducted to examine cross-sectional association of each metabolite with baseline diagnosis (6 models per metabolite). We performed logistic regression models to compare BA levels between the MCI-NonConverter and MCI-Converter groups. Cox proportional hazard models were used to evaluate the association of metabolite levels with progression from MCI (combined EMCI and LMCI subjects) to AD. The cross-sectional association of ADAS-Cog13 with BAs was assessed using linear regression models with square root of ADS-Cog13 as the dependent variable.

In ROS/MAP, one sample per individual was used. Linear regression models with global cognition score as dependent variable and metabolites as independent variables were used to assess the association of serum BAs with cognition, while adjusting for sex, age, *APOE* ε4, and years of education. Similar analyses were conducted for brain BAs separately.

We restricted our genetic variant analysis to single nucleotide polymorphisms (SNPs) in genes involved in immune response pathway that were significantly associated with AD genome-wide[2, 56-58]. Selected genetic variant included rs616338-T(*ABI3*), rs143332484-T(*TREM2*), rs72824905-C(*PLCG2*), rs9331896-T(*CLU*), rs6656401-A(*CR1*), rs35349669-T(*INPP5D*), rs11771145-G(*EPHA1*), rs983392-A(*MS4A6A*), and rs190982 -A(*MEF2C*). Associations of AD risk variants in immune-related genes with selected metabolic traits in ADNI and RS were computed using sex, age, and BMI as covariates.

## 3. Results

Characteristics of ADNI participants are depicted in **Table 1**. Baseline cognitive measurements were significantly different among diagnostic groups, as expected. AD patients were more often carriers of at least one *APOE* ε4 allele. In addition, ADAS-Cog13 scores were not significantly different between the MCI-converter and NonConverter groups. However, the proportion of *APOE* ε4 carriers was higher in MCI-Converter group.

### 3.1. Serum BA profiles are significantly altered in AD

The Bonferroni-corrected threshold for statistical significance was determined as *P* < 4.76 × 10^−4^ (0.05 divided by 15 metabolites times 7 phenotypes including cognition). When we compared BA profile in AD to CN, we detected a significant decrease in levels of the primary BA, CA (*P =* 1.56 × 10^−4^). In contrast, a significant increase of bacterially produced secondary BA, DCA was noted (*P =* 1.61 × 10^−4^) along with several secondary conjugated BAs, GDCA, TDCA, and GLCA (**Table 2**). GDCA and GLCA were significantly associated with ADAS-Cog13 where higher levels indicate worse cognition. Comparing BA levels between AD and both MCI groups yielded similar results, while the comparison of BA levels between the CN and MCI groups did not reach statistical significance (**Supplementary Table 4**).

### 3.2. Ratios reflective of conversion of BAs by gut microbiome are significantly associated with AD and cognitive performance

To determine which enzymatic processes in BA metabolism may underlie the differences noted in AD, we investigated eight selected ratios reflective of enzymatic activities in the liver and the gut microbiome. These ratios included:

1. The CA:CDCA ratio was selected to test if a possible shift in BA synthesis from the primary to the alternative BA pathway occurs in the liver.
2. Ratios of secondary to primary BAs (DCA:CA, GLCA:CDCA, and TLCA:CDCA) to investigate differences in gut microbiome enzymatic activity leading to altered production of secondary BAs. Since LCA was excluded in QC steps, the GLCA:CDCA and TLCA:CDCA ratios were used as proxies for LCA:CDCA ratio.
3. GDCA:DCA and TDCA:DCA ratios were used to test if the observed secondary BA dysregulation is related to enzymatic differences related to their taurine and glycine conjugation.

Here, we considered associations as significant at a Bonferroni-corrected *P* < 3.11 × 10^−4^ (0.05 divided by all 23 metabolic traits times 7 phenotypes, which include cognition). The ratio of the primary BAs (CA:CDCA) showed no significant association with AD. Yet, for the ratio of DCA:CA, i.e. the conversion of unconjugated primary to unconjugated secondary BA, we observed a highly significant association with AD diagnosis (*P=*1.53 × 10^−8^). Ratios between primary and secondary conjugated BAs showed the same effect and direction, including GDCA:CA (*P=*8.53 × 10^−10^), TDCA:CA (*P=*9.83 × 10^−7^), and GLCA:CDCA (*P=*3.61 × 10^−6^). Ratios modeling the glycine and taurine conjugation step of DCA, i.e. GDCA:DCA, TDCA:DCA, were not significantly associated with diagnosis (**Fig. 2** and **Table 3**).

**Table 3.** Ratios of bile acids reflective of gut microbiome and liver enzymatic activities and their correlation with disease status and cognitive function^a^.

| **Ratios informative about metabolic processes** | **Ratios calculated** | **CN vs. AD (*n*=673) OR (95% CI); *P*-value<sup>b</sup>** | **ADAS-Cog13 (*n*=1453) $\beta$(95% CI); *P*-value<sup>c</sup>** |
| --- | --- | --- | --- |
| Bile acid synthesis: primary vs. alternative pathway | CA:CDCA | 0.87(0.77,0.97);1.67E-02 | -0.03(-0.07,0.01);1.27E-01 |
| Conversion from primary to secondary BA by the gut microbiome | DCA:CA | **1.25(1.16,1.35);1.53E-08** | **0.05(0.03,0.08);1.05E-05** |
|  | GDCA:CA | **1.24(1.16,1.33);8.53E-10** | **0.06(0.04,0.08);1.20E-07** |
|  | TDCA:CA | **1.16(1.10,1.24);9.83E-07** | **0.04(0.02,0.06);5.40E-05** |
|  | GLCA:CDCA | **1.16(1.09,1.23);3.61E-06** | **0.04(0.02,0.06);9.15E-05** |
|  | TLCA:CDCA | 1.09(1.03,1.16);1.60E-03 | 0.03(0.01,0.05);1.50E-03 |
| Glycine or Taurine conjugation of secondary bile acids by liver enzymes | GDCA:DCA | 1.16(1.02,1.31);2.41E-02 | 0.05(0.02,0.10);5.49E-03 |
|  | TDCA:DCA | 1.02(0.93,1.11);7.40E-01 | 0.01(-0.02,0.04);4.15E-01 |

<sup>a</sup>Several ratios were calculated to inform about possible enzymatic activity changes in Alzheimer's patients. These ratios reflect: (1) Shift in bile acid metabolism from primary to alternative pathway. (2) Changes in gut microbiome correlated with production of secondary bile acids. (3) Changes in glycine and taurine conjugation of secondary bile acids.
<sup>b</sup>Outcome: Baseline diagnosis; Odds ratios and p-values were obtained from logistics regressions. Models were corrected for age, sex, body mass index, and APOE ε4 status; Bonferroni-adjusted critical value was set to 1.04E-03 based on 6 possible pairwise comparison of diagnosis groups (CN, EMCI, LMCI, and AD) for 8 ratios.
<sup>c</sup>Outcome: Square root of ADASCog-13 (0 [best] to 85 [worst]); Models were corrected for age, sex, years of education, body mass index, and APOE ε4 status; Bonferroni-adjusted critical value was set to $.11 \times 10^{-4}$ (0.05 divided by all 23 metabolic traits times 7 phenotypes, which include cognitive function).

**Fig. 2.**
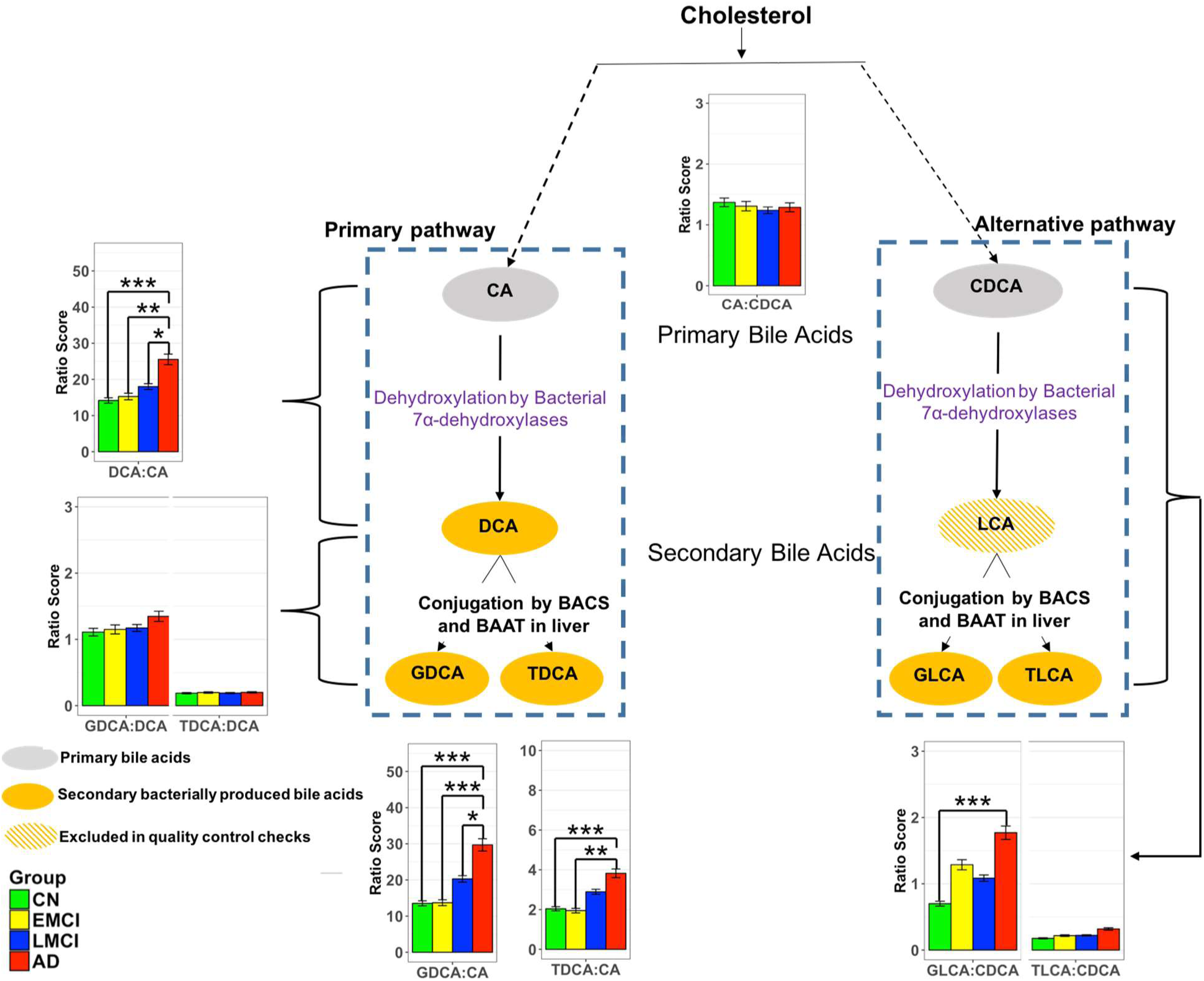
Ratios of bile acids reflective of liver and gut microbiome enzymatic activities in CN, Early MCI, Late MCI and AD patients. Three types of ratios were calculated to inform about possible enzymatic activity changes in Alzheimer’s patients. These ratios reflect one of the following: (1) Shift in bile acid metabolism from primary to alternative pathway. (2) Changes in gut microbiome correlated with production of secondary bile acids. (3) Changes in glycine and taurine conjugation of secondary bile acids. Color code: Green: cognitively normal; Yellow: EMCI; Blue: LMCI; Red: AD. Composition of selected ratios stratified by clinical diagnosis. Error bars indicate standard error of the means; Asterisks indicate statistical significance (\**P*<10^−03^, ** *P*< 10^−04^, and \*\*\**P*< 10^−05^). *P*-values were estimated from logistic regression models and adjusted for age, sex, body mass index, and *APOE* ε4 status. The significance level was adjusted for multiple testing according to Bonferroni method to 0.05/138 = 3.62E-4; LCA were excluded in the quality control steps.

Four ratios (including DCA:CA and GLCA:CDCA) were significantly associated with ADAS-Cog13. For the ratios we observed the same pattern as for AD diagnosis, with higher ratios of secondary to primary BAs being highly significantly associated with worse cognitive performance, while neither conjugation, nor a shift between primary and alternative BA pathways in the liver were significantly linked to cognition (**Table 2** and **3**).

### 3.3. Serum BA levels were associated with progression from MCI to AD in ADNI

The 9 metabolites and ratios associated with diagnosis were further investigated to assess their relationship with progression from MCI to AD. Out of 779 MCI (EMCI and LMCI) patients with mean (SD) follow-up 3.94 (2.35), 32.2% progressed to AD dementia in four years (labeled as MCI-Converter (*n*=251) vs. those that did not progressed MCI-NonConverter (*n*=528)). BA profiles were compared between the two groups using logistic regression models with conversion status as dependent variable and metabolite as independent variable. Models were adjusted for age, sex, BMI, baseline ADAS-Cog13 score, and *APOE* ε4. The Bonferroni-corrected threshold for statistical significance was determined as *P* < 5.56 × 10^−3^ (0.05 divided by 9 metabolites and ratios). We noted a decrease in CA levels (*P=*9.12 × 10^−4^) and an increase in ratios of GDCA:CA (*P=*1.63 × 10^−3^) and TDCA:CA (*P=*1.72 × 10^−3^) in MCI-Converters (**Fig. 3** and **Supplementary Table 5**). Further survival analysis also revealed that levels of CA (hazard ratio (HR), 0.92; *P* =3.79 × 10^−3^), GDCA:CA (HR, 1.07; *P=*2.81 × 10^−3^), and TDCA:CA (HR, 1.06; *P=*3.19 × 10^−3^) ratios predicted MCI progression (**Fig. 3**).

**Fig. 3.**
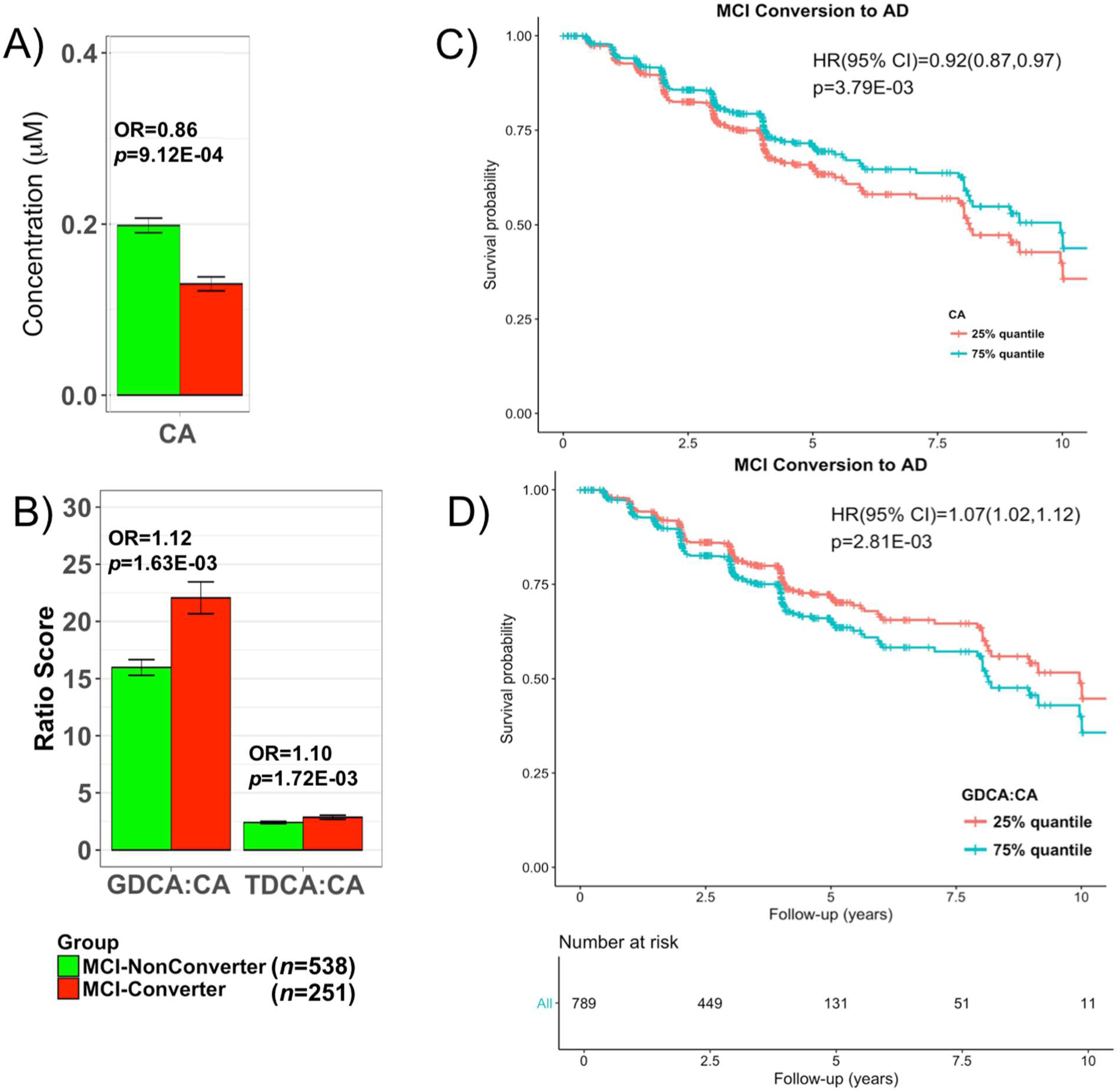
Comparison of bile levels in MCI subjects who convert and those who did not convert to AD dementia. **A** and **B**. Lower levels of CA and higher levels of two secondary to primary ratios were significantly associated with higher odds of converting from MCI to AD. EMCI and LMCI patients that converted to AD dementia in 4 years after baseline were labeled as MCI-Converter; 9 bile acids and ratios that were significantly dysregulated between CN to AD were assessed; *P*-values were estimated from logistic regression models and adjusted for age, sex, body mass index, and *APOE* ε4 status; the significance level was adjusted for multiple testing according to Bonferroni 0.05/9 = 5.56 × 10^−3^. **C** and **D**. Cox hazards model of the association of conversion from MCI to AD. Red line: 1st quantile, Red line: 3rd quantile. Analysis was conducted using quantitative values and stratification by quantiles was used only for graphical representation.

### 3.4. Replication of association between cognition and DCA:CA ratio in serum and brain from ROS/MAP

In order to confirm the associations observed in ADNI, we used an independent cohort of older adults (ROS/MAP) with measures of BAs in serum and brain to replicate our findings. Since the sample sizes in ROS/MAP were smaller than ADNI and AD cases were strongly underrepresented (*n*=11 for the serum samples), we focused on replicating the association between global cognition (where higher values indicate better cognition, which is in contrast to ADNI where higher scores indicate worse cognition) and the DCA:CA ratio (as proxy for BA processing by the gut microbiome). Separate linear regression models were used for brain and serum samples. Pearson’s correlation coefficient between serum DCA:CA and DCA:CA in 93 matching brain samples was 0.303 (*P*=0.003). In both serum and brain samples, higher levels of DCA:CA were associated with worse cognition (serum: β = −0.06; *P =* 0.011; brain: β = −0.21; *P =* 0.032), replicating our ADNI finding.

### 3.5. Genetic risk variants for AD in genes related to immune function are associated with bile acid levels

To further evaluate that altered BA profiles in AD are related to processes in the gut microbiome, we investigated if BA profiles were associated with immune-related AD risk genes which may contribute to differences in gut microbiome composition. Using the ADNI (*n*=817 with WGS data) and RS (*n*=488) cohorts, association of selected BAs in the primary BA pathway (CA, DCA, GDCA, and TDCA) as well as the DCA:CA ratio with the selected genetic risk variants in 9 candidate genes with immune-related functions was assessed. In addition, we included associations from a published large cohort-based study[44] to increase sample size. With the exception of rs983392 in *MS4A6A*, we found nominally significant associations for the candidate variants in all of these genes (**Supplementary Table 10**). Three associations were significant after Bonferroni-correction (*P* < 1.1 × 10^−3^) in at least one of the studies: rs616338 (*ABI3*) and rs190982 (*MEF2C*) were significantly associated with the DCA:CA ratio, and rs11771145 (*EPHA1*) was significantly linked to both DCA and TDCA.

### 3.6. Genetic loci associated with DCA may influence susceptibility for AD

To follow up on the hypothesis that elevated DCA levels in AD linked to gut dysbiosis are relevant in the pathogenesis of AD, we collected (suggestive) significant genetic associations with DCA levels (*P <* 1.0 × 10^−5^) from a previous study of genetic influences on blood metabolite levels in large population-based cohorts (n∼7,800)[44]. We then annotated the resulting 13 loci with genetic trait associations, including AD associations from the IGAP study[2], and tried to replicate associations with DCA in ADNI (**Supplementary Table 11**). Two of the 13 genes, *CYP7A1* and *IMPA2*, also showed association with DCA levels in ADNI subjects. Notably, six of the 13 genes have been previously linked via genetic studies to AD (*ABCA7*) or AD phenotypes, including cognitive decline and CSF protein levels (*LRRC7, CYCS, GPC6, FOXN3* and *CNTNAP4*).

## 4. Discussion

Cholesterol metabolism has been extensively implicated in the pathogenesis of AD through biological, epidemiological, and genetic studies, yet the molecular mechanisms linking cholesterol and AD pathology are still not well understood. Many cholesterol metabolism related genes such as *BIN1, CLU, PICALM, ABCA7, ABCG1*, and *SORL1* are classified among the top 20 late onset AD susceptibility loci by some of the largest genome-wide association studies (GWAS) undertaken to date[2, 19]. The ε4 allele of the apolipoprotein gene[59, 60], the most robust and reproducible genetic risk factor for AD, is involved in the transport of cholesterol.

In this study, we interrogated a possible role for BA end products of cholesterol metabolism and clearance in cognitive changes in AD. BA are produced in liver and by the gut microbiome, provide key regulatory functions in energy homeostatic mechanisms, and are indicators of gut dysbiosis. Using stored blood samples from ADNI studies we established that BA profile is significantly altered in AD patients with changes detected earlier in disease (**Fig. 1** and **2**). We noted a significant decrease in serum levels of a liver-derived primary BA (CA) and an increase in levels of a bacterially produced secondary BAs and their conjugated forms (DCA, GDCA and TDCA, GLCA) in AD patients compared to cognitively normal subjects (**Table 2, Fig. 1A**). Higher levels of secondary conjugated BAs (GDCA, GLCA, and TLCA) were significantly associated with worse cognitive function measured by the ADAS-Cog13 (**Table 2, Fig. 1A**).

To inform about enzymatic activity changes in liver and gut, three types of metabolite ratios were evaluated to inform about mechanisms leading to the noted altered BA profile in AD. We found no shift in metabolism between primary and alternative pathways (**Fig. 2**; no change in CA:CDCA); a significant change in production of secondary BAs via enzymatic activities in gut microbiome (increased DCA:CA as well as GLCA:CDCA and TLCA:CDCA as proxies for LCA:CDCA) and no change in processes involved in glycine and taurine conjugation of secondary BAs in the liver (no change in GDCA:DCA and TDCA:DCA). The significant increase in ratios of secondary to primary BAs (e.g. DCA:CA; **Fig. 2**), suggest altered activity of bacterial 7α-dehydroxylases leading to excess production of secondary BAs many of which were previously reported as cytotoxic[34, 61-63]. This indicates potential gut dysbiosis in AD patients possibly caused by enhanced colonization of the large and possibly the small intestine with anaerobic bacteria capable of CA and CDCA 7α-dehydroxylation. Increases in these ratios also significantly correlated with poorer cognition (**Table 2**). Together, these findings suggest that enzymatic steps in conversion of primary to secondary BAs in the gut might contribute to disease.

We also evaluated effects of BA levels on risk of progression to AD among 538 MCI patients. We noted that lower levels of CA and higher ratio of secondary to primary BAs, GDCA:CA, and TDCA:CA were significantly associated with risk of developing AD dementia (**Fig. 3, Supplementary Table 6**).

The increased production of bacterially produced DCA from CA modeled by ratio DCA:CA and its link to cognition was replicated in the independent ROS/MAP cohort. Association of the DCA:CA ratio with disease severity was evaluated separately in serum and brain samples. Due to the small number of AD patients (*n*=11), we used global cognitive score as an index of disease severity. Similar to ADNI findings, an increase in the DCA:CA in both serum and brain were significantly associated with worse cognitive outcome. This finding suggests that downstream effects of the gut-directed dysregulation of primary vs. secondary BAs are not limited to the periphery, but also might affect metabolic homeostasis and/or signaling functions in the human brain.

Earlier smaller studies suggested differences in BA levels in AD[26, 38-40]. For example, in a study of 495 plasma metabolites comparing metabolite levels among MCI (*n*=58) and AD (*n* = 100) with those of cognitively normal controls (*n*=93), levels of DCA, LCA, and GLCA were significantly elevated in the disease state[40]. Mapstone and colleagues[38] identified increased levels of glycoursodeoxycholic acid (GUDCA) in subjects likely to develop amnestic MCI or AD within the next 2 to 3 years compared to the control group. We replicated these findings with the exception of LCA that was excluded during QC and GUDCA, which showed only a non-significant trend of upregulation in the AD group (*P = 0.0*54).

Composition and functional changes of the gut microbiome have been implicated in several diseases. Microbiome GWAS reveal that variants in many human genes involved in immunity and gut architecture are associated with an altered composition of the gut microbiome[64]. Although many factors such as diet can affect the microbial organisms residing in the gut, emerging data support the hypothesis that certain host genetic variants predispose an individual towards microbiome dysbiosis and this can be linked to disorders of metabolism and immunity such as Crohn’s disease, ulcerative colitis, type 2 diabetes mellitus, asthma, obesity, autism and rheumatoid arthritis[64].

Accumulating evidence links dysregulation of the immune system to AD pathology. In particular, genetic association studies in AD have robustly identified several genetic risk variants in immune-related genes[2, 58]. Using the ADNI and RS cohorts, we investigated the association of BA profiles of CN subjects with genetic variants in nine AD-related and innate immunity genes. Eight of them were associated with selected BA levels at nominal significance (**Supplementary Table 10**). Three of these associations were significant after Bonferroni-correction, with rs616338 (*ABI3*) and rs190982 (*MEF2C*) associated with the DCA:CA ratio, and rs11771145 (*EPHA1*) linked to both DCA and TDCA. The association of the BAs to AD genes suggest that these immune related genes may influence the risk of AD through the BA metabolism or changes in the gut microbiome. Interestingly, both *ABI3* and *MEF2C* are thought to be involved in immune reactions to pro-inflammatory stimuli that are partially secreted by microbes[65, 66]. The link to the DCA:CA ratio may thus mirror differences in gut microbiome composition due to altered immune response in AD, providing a mechanistic hypothesis for our findings. The function of *EPHA1* is not well understood but, it has been hypothesized that when activated, this receptor may affect the integrity of the blood brain barrier (BBB)[67]. Its association with levels of DCA is intriguing as DCA is known to be cytotoxic and can disrupt the BBB and then enter the brain[28]. rs11771145 is associated with gene expression levels of *EPHA1*[54], and as DCA is not known to be produced by human metabolism, changed expression and activity of *EPHA1* may be related to DCA-mediated cytotoxic effects.

Using an established atlas of genetic influences on human blood metabolites[44], we further investigated a potential cytotoxic role of DCA. For almost half of the 13 identified loci, we found genetic evidence for involvement in AD-linked complex traits (**Supplementary Table 11**). In particular, *ABCA7* is an AD risk gene replicated in several genetic studies[68, 69]. Five additional genes (*LRRC7, CYCS, GPC6, FOXN3*, and *CNTNAP4*) genetically influence AD phenotypes, including cognitive decline and CSF markers. While it remains speculative if and how these genes interact with DCA to contribute to AD risk, it is intriguing that we identified *ABCA7* by screening for associations with DCA levels. *ABCA7* is highly expressed in the brain, and functions in the efflux of lipids, including cholesterol, from cells. Due to the structural similarity of DCA and cholesterol, we hypothesize that *ABCA7* may be able to also transport this BA, reconciling metabolomics findings via a functional hypothesis to a risk gene for AD. The findings that BA levels are regulated by AD related genes might provide new mechanistic insights.

There is growing support for strong connections between the intestinal environment, with its diverse microbial composition and activity, and the functions of the central nervous system. The “gut-brain metabolic axis” facilitates bidirectional chemical communication between the central and enteric nervous systems through mechanisms just starting to be defined[7-9]. Such a metabolic axis is thought to be involved in the regulation of multiple host metabolic pathways in which levels of hormones, neurotransmitters, amines, GABA, short-chain fatty acids (SCFA), lipid metabolites, and others are regulated by gut microbiome activity[12]. Changes in the composition of intestinal bacterial populations are associated with a wide array of conditions, including neurological and neurodevelopmental disorders such as multiple sclerosis, autism, depression, schizophrenia, and Parkinson’s disease[70-72]. In addition, increasing evidence suggest that liver disease may impact cognitive functions and contribute to AD[73].

Our findings suggest novel metabolic links in AD where BAs represent a component of the gut-liver-brain axis that relates to cognitive functions. It is of interest that BAs that are ligands for the nuclear transcription factor FXR, which along with other nuclear receptors acts synergistically as metabolic sensors to regulate energy homeostasis pathways[74, 75] peripherally might also propagate their effects to the brain. Interestingly, levels of four BAs that are produced by the gut microbiome and that we show to be significantly correlated with disease status and cognition (DCA, GLCA, TLCA, TDCA) are hydrophobic and cytotoxic[34, 35, 76, 77]. Cell lines, animal models, and human studies suggest that levels of such BA, particularly DCA, lead to a disruption of mitochondrial membranes resulting in increased reactive oxygen species, markers of inflammation, and apoptosis as well as decreases in cell viability and DNA synthesis[34, 35, 78]. DCA increases BBB permeability with deuterium labelled DCA showing it crosses the BBB in rodents[27, 29]. Increased amounts of secondary BAs in blood may enter the brain through induced permeability of the BBB, affecting brain physiology and metabolism[28]. Additionally, the BA nuclear receptor FXR, TGR5, and transporters Ostα/Ostβ seem to be expressed in the brain[79]. Several studies in human and animal brains also revealed that the full panel of BAs are found in the brain[24-27], but it is unclear whether this is due to transport from the periphery, from local synthesis, or both. The function of these BAs in the brain remains poorly defined with some support for them acting as neurosteroids[80]

BA levels and the gut microbiome influence each other, where BSH-rich bacteria readily modify the BA profile while, on the other hand, intestinal BAs control the growth and maintenance of commensal bacteria, maintain barrier integrity, and modulate the immune system[81-84]. Such changes might impact brain functions. Future longitudinal studies covering pre-symptomatic stages are needed to establish the influence of immune changes on gut microbiome composition and activity in AD patient. Tracking earliest changes in BA and other gut derived metabolites might provide insights into causality. Labeling studies are needed to evaluate if BAs cross the BBB and build up in brain with further elucidation of their signaling and regulatory functions centrally. However, we cannot exclude the possibility that changes in the brain during disease can also impact the gut and liver, and hence some of our findings might be brain derived.

### 4.1. Limitations

This is an observational study, the results of which may contain confounding biases. For example, diet, lifestyle, exposome and other factors may contribute to changes in the gut. It remains unclear how these important factors are related to AD pathogenesis and whether the observed differences we note are causes or consequences of disease. Further studies of metabolic changes in normal aging are required to help define which aspects of BA metabolism might be related to disease vs normal aging. Use of medications was extensively evaluated as a possible confound (**Supplementary methods and Tables 8-10**) and our key findings remained after controlling for medication use but larger studies need to further evaluate the effect of these medications. Additional experimental studies are needed to more fully define the expression of BAs and their receptors in the brain and mechanistic roles of BAs in the development of AD. The impact of BAs on FXR, TGR5, vitamin, and hormone receptors in the brain and the signaling pathways impacted are currently unclear. It is important to evaluate in other large community studies the generalizability of our findings. The genetic links need to be tested in large populations.

## 5. Conclusions

In summary, there is evidence of a relationship among intestinal BA profile, gut microbial composition and/or activity, innate immunity, and genetic variants implicated in AD. When disrupted, BAs may contribute to cognitive changes, highlighting the importance of cholesterol clearance and its regulation in AD. Disorders in BA metabolism cause cholestatic liver diseases, dyslipidemia, fatty liver diseases, cardiovascular diseases, and diabetes, which are all associated with risk of cognitive decline, directly or indirectly. Our results lend support to this relationship in the context of AD and cohorts at risk for AD. Our evolving understanding of the gut microbiome’s role in aging and in central nervous system diseases and their progression could open potential new hypotheses in the field, regardless of whether the role is ultimately found to be causative, consequence, or contributory. The role of the gut microbiome in AD needs to be further investigated along with the emerging links between central and peripheral metabolic failures that might contribute to brain health and disease during aging.

## Author Contributions

MahmoudianDehkordi, Arnold and Nho had full access to all of the data in the study and take responsibility for the integrity of the data and the accuracy of the data analysis.

**Statistical analyses also included:** Toledo, Arnold, Bhattacharyya, Jia, Massaro, Rotroff, Jack, Kastenmüller

**Data management and medication term mapping:** Tenenbaum and Blach

**Concept and design:** Kaddurah-Daouk lead concept and design team that included all co-authors

**Data analysis and quality control of metabolomic data:** Schafferer, Klein, Koal, St. John Williams, Thompson, Moseley

**Acquisition, quality control and processing of metabolomic data:** St. John Williams, Thompson, Moseley

**Drafting of the manuscript:** MahmoudianDehkordi, Nho, Arnold, Louie, Kastenmüller, Kueider-Paisley, Kaddurah-Daouk

**Biochemical, genomics and medications integration:** Kastenmüller, Baillie, Han, Risacher, Arnold; Nho

**Data deposition:** Alzheimer’s Disease Neuroimaging Initiative (see note)

**Harmonization of methods:** Alzheimer’s Disease Metabolomics Consortium

**Technical, bibliographic research and/or material support:** Louie

**Biochemical interpretation:** Baillie, Han, Kaddurah-Daouk, Jia, Bhattacharyya, Arnold

**Critical revision of the manuscript for important intellectual content:** Saykin, Doraiswamy, Kastenmüller, Kaddurah-Daouk

**Obtained funding:** Kaddurah-Daouk

**Supervision:** Motsinger-Reif, Trojanowski, Shaw, Weiner, Doraiswamy, Saykin, Kastenmüller, Kaddurah-Daouk

**The Alzheimer’s Disease Metabolomics Consortium (ADMC):** A complete listing of ADMC investigators can be found at: https://sites.duke.edu/adnimetab/who-we-are/

**The Alzheimer’s Disease Neuroimaging Initiative (ADNI):** Data used in the preparation of this article were obtained from the ADNI database (http://adni.loni.usc.edu). As such, the investigators within the ADNI contributed to the design and implementation of ADNI and/or provided data but did not participate in analysis or writing of this report. A complete listing of ADNI investigators can be found at: http://adni.loni.usc.edu/wp-content/uploads/how_to_apply/ADNI_Acknowledgement_List.pdf.

## Funding/Support

Funding for ADMC (Alzheimer’s Disease Metabolomics Consortium, led by Dr R.K.-D. at Duke University) was provided by the National Institute on Aging grant R01AG046171, a component of the <u>Accelerated Medicines Partnership for AD (AMP-AD) Target Discovery and Preclinical Validation Project</u> (https://www.nia.nih.gov/research/dn/amp-ad-target-discovery-and-preclinical-validation-project) and the National Institute on Aging grant RF1 AG0151550, a component of the M^2^OVE-AD Consortium (Molecular Mechanisms of the Vascular Etiology of AD – Consortium https://www.nia.nih.gov/news/decoding-molecular-ties-between-vascular-disease-and-alzheimers).

Data collection and sharing for this project was funded by the Alzheimer’s Disease Neuroimaging Initiative (A.D.N.I.) (National Institutes of Health Grant U01 AG024904) and DOD A.D.N.I. (Department of Defense award number W81XWH-12-2-0012). A.D.N.I. is funded by the National Institute on Aging, the National Institute of Biomedical Imaging and Bioengineering, and through generous contributions from the following: AbbVie, Alzheimer’s Association; Alzheimer’s Drug Discovery Foundation; Araclon Biotech; BioClinica, Inc.; Biogen; Bristol-Myers Squibb Company; CereSpir, Inc.; Eisai Inc.; Elan Pharmaceuticals, Inc.; Eli Lilly and Company; EuroImmun; F. Hoffmann-La Roche Ltd and its affiliated company Genentech, Inc.; Fujirebio; GE Healthcare; IXICO Ltd.; Janssen Alzheimer Immunotherapy Research & Development, LLC.; Johnson & Johnson Pharmaceutical Research & Development LLC.; Lumosity; Lundbeck; Merck & Co., Inc.; Meso Scale Diagnostics, LLC.; NeuroRx Research; Neurotrack Technologies; Novartis Pharmaceuticals Corporation; Pfizer Inc.; Piramal Imaging; Servier; Takeda Pharmaceutical Company; and Transition Therapeutics. The Canadian Institutes of Health Research is providing funds to support A.D.N.I. clinical sites in Canada. Private sector contributions are facilitated by the Foundation for the National Institutes of Health (www.fnih.org). The grantee organization is the Northern California Institute for Research and Education, and the study is coordinated by the Alzheimer’s Disease Cooperative Study at the University of California, San Diego. A.D.N.I. data are disseminated by the Laboratory for Neuro Imaging at the University of Southern California.

The work of various Consortium Investigators are also supported by various NIA grants [U01AG024904-09S4, P50NS053488, R01AG19771, P30AG10133, P30AG10124, K01AG049050], the National Library of Medicine [R01LM011360, R00LM011384], and the National Institute of Biomedical Imaging and Bioengineering [R01EB022574]. Additional support came from Helmholtz Zentrum, the Alzheimer’s Association, the Indiana Clinical and Translational Science Institute, and the Indiana University-IU Health Strategic Neuroscience Research Initiative.

### Role of the Funder/Sponsor

[Funders listed above] had no role in the design and conduct of the study; collection, management, analysis, and interpretation of the data; preparation, review, or approval of the manuscript; and decision to submit the manuscript for publication.

### Additional Contributions

The authors are grateful to Lisa Howerton for administrative support and the numerous ADNI study volunteers and their families.

## DISCLOSURES

J.B.T. reports investigator-initiated research support from Eli Lilly unrelated to the work reported here. S.S., S.K, and T.K. are employed by Biocrates Life Sciences AG. These authors have no other financial relationships relevant to this article to disclose. J.Q.T. may accrue revenue in the future on patents submitted by the University of Pennsylvania wherein he is a co-inventor and he received revenue from the sale of Avid to Eli Lily as a co-inventor on imaging-related patents submitted by the University of Pennsylvania. L.M.S. receives research funding from NIH (U01 AG024904; R01 MH 098260; R01 AG 046171; 1RF AG 051550); MJFox Foundation for PD Research and is a consultant for Eli Lilly; Novartis; Roche; he provides QC over-sight for the Roche Elecsys immunoassay as part of responsibilities for the ADNI3 study. A.J.S. reports investigator-initiated research support from Eli Lilly unrelated to the work reported here. He has received consulting fees and travel expenses from Eli Lilly and Siemens Healthcare and is a consultant to Arkley BioTek. He also receives support from Springer publishing as an editor in chief of Brain Imaging and Behavior. M.W.W. reports stock/stock options from Elan, Synarc, travel expenses from Novartis, Tohoku University, Fundacio Ace, Travel eDreams, MCI Group, NSAS, Danone Trading, ANT Congress, NeuroVigil, CHRU-Hopital Roger Salengro, Siemens, AstraZeneca, Geneva University Hospitals, Lilly, University of California, San Diego–ADNI, Paris University, Institut Catala de Neurociencies Aplicades, University of New Mexico School of Medicine, Ipsen, Clinical Trials on Alzheimer’s Disease, Pfizer, AD PD meeting. PMD has received research grants and advisory/speaking fees from several companies for other projects, and he owns stock in several companies. Full disclosures will be made through the IJCME form. R.K.D. is inventor on key patents in the field of metabolomics including applications for Alzheimer disease. All other authors report no disclosures.

An email with links to the Authorship Form will be sent to authors for completion after manuscripts have been submitted

## Abbreviations

AD: Alzheimer’s disease;
ADAS-Cog13: 13-item Alzheimer’s Disease Assessment Scale-Cognitive subscale;
ADMC: Alzheimer’s Disease Metabolomics Consortium;
ADNI: Alzheimer’s Disease Neuroimaging Initiative;
APOE: apolipoprotein E;
ASBT: Apical sodium-dependent bile acid transporters;
BBB: blood-brain barrier;
BSEP: Bile salt export pump;
BSH: Bile salt hydrolases;
BA: bile acid;
CA: Cholic acid;
cAMP: Cyclic adenosine monophosphate;
CDCA: Chenodeoxycholic acid;
CN: Cognitively normal older control;
DCA: Deoxycholic acid;
EMCI: Early mild cognitive impairment;
FDR: False discovery rate;
FXR: Farnesoid X Receptor;
GCA: Glycocholic acid;
GCDCA: Glycochenodeoxycholic acid;
GDCA: Glycodeoxycholic acid;
GLCA: Glycolithocholic acid;
GUDCA: Glycoursodeoxycholic acid;
LCA: Lithocholic acid;
LMCI: Late mild cognitive impairment;
MCI: Mild cognitive impairment;
NTCP: Sodium/Taurocholate co-transporting polypeptide;
OST-α: Organic solute transporter alpha;
OST-β: Organic solute transporter beta;
ROS/MAP: The Religious Orders Study (ROS) and the Memory and Aging Project (MAP);
RS: Rotterdam Study;
SHP: Small heterodimer partner;
SMC: Subjective memory concern;
TCA: Taurocholic acid;
TCDCA: taurochenodeoxycholic acid;
TDCA: Taurodeoxycholic acid;
TLCA: Taurolithocholic acid;
TMCA: Trimethoxycinnamic acid;
TUDCA: Tauroursodeoxycholic acid;
UDCA: Ursodeoxycholic acid

