## Supplementary Materials for "Altered Bile Acid Profile Associates with Cognitive Impairment in Alzheimer’s Disease – An Emerging Role for Gut Microbiome"

**Supplementary Materials and Methods**

### 1. ADNI bile acid profiles: Quality control and medication adjustment

#### 1.1. Quality control of ADNI bile acid profiles

The preprocessing included the steps briefly described herein**.** In summary, two sulphates (Glycolithocholic acid sulfate and Taurolithocholic acid sulfate) were excluded from dataset as Biocrates has not analytically validated the results for these compounds. Metabolites with >40% of measurements below the lower limit of detection (<LOD) were excluded. To assess the precision of the measured analytes, a set of blinded analytical replicates (93 samples for 24 individuals) were supplied by the Alzheimer’s Disease Neuroimaging Initiative (ADNI); keeping the Metabolomics Consortium blinded of the matching pairs. Following unblinding the bile acid (BA) profiles, the accuracy was measured using the intra-class correlation coefficient (ICC) and coefficient of variation (CV) using the blinded replicates. Using blinded replicated samples, metabolites with CV>25% across plates, and ICC <0.65 were excluded from subsequent analyses. These three steps, reduced the total number of metabolites to 15 (**Table 2**). <LOD values were imputed using LOD values divided by 2 for each specific metabolite. Non-fasting participants were flagged for removal from the datasets. For the participants with replicated measurements, the average of the measurements was computed to give one value per biological sample in further analyses. The presence of multivariate outliers was examined by evaluating the first and second principle components. The datasets were extended by 8 ratios of BAs. Log_2_ transformation was performed for all metabolites and ratios.

After performing the quality control steps outlined above, the final ADNI dataset comprised 1464 individuals (average age (SD). = 73.62 (7.21); 808 men/656 women) including 305 Alzheimer’s disease (AD), 505 Late mild cognitive impairment (LMCI), 284 Early mild cognitive impairment (EMCI), and 370 healthy participants. All samples retained were taken with the participant in a fasting state.

#### 1.2. Medication Adjustment of ADNI bile acid profiles

The preprocessed BA values obtained from the quality control step was used for subsequent association analyses was adjusted to take into account the effect of medications on the levels of the BAs. In ADNI, 42 major medication classes used to treat psychiatric (including different categories of benzodiazepines, antipsychotics, and antidepressants) and cardiovascular conditions (including different categories of anti-hypertensives, cholesterol treatment, and anti-diabetics), as well as dietary supplements (Co-Q10, fish oil, nicotinic acid, and acetyl L-carnitine) were systematically coded and available for model-based evaluations of the influence of each drug type on metabolite levels. Intake of any medication within a category was coded as present or absent. Two AD medication classes, i.e. cholinesterase inhibitors and memantine, were excluded from this process due to high collinearity (Cramer’s V = 0.65 and 0.52, respectively) of these medications with AD diagnosis.

Medication information, including dosage and reason for taking, was collected from research participants in the form of free text. In order to account for exposure to specific drugs and drug classes in our analysis, it was necessary to convert that unstructured data into structured terms representing both the specific drug and the drug class[es] to which it belonged. We used the RxNorm API (application programming interface) to convert drug names, including synonyms and misspellings, into coded drug terms from RxNorm, a standardized drug terminology developed and maintained by the National Library of Medicine (NLM). Inexact matches were manually reviewed and corrected where necessary. The RxNorm API was then used to identify corresponding drug classes for the respective coded medications. For example, citalopram, citalopran, citalporam, and Celexa all mapped to the concept “citalopram” with RxNorm Concept Unique Identifier “2556”. Along with drugs like Zoloft, Lexapro, and Prozac, they were mapped to the classes “Antidepressive Agents, Second-Generation” and “Serotonin Reuptake Inhibitor.” An iterative approach was used to identify drug classes of interest from hundreds of partially overlapping possible classifications. Drug classes were identified for a core set of medications.  The other medications sharing these classes were determined and all of their respective drug classes were identified, further generating new medications and classes. Through iteration and pruning based on review with clinical experts, the final set of ontology classes of interest was created.

AD medications (Cholinesterase Inhibitor and N-methyl-D-aspartate receptor antagonist) are a special issue in this context. Because these medications are taken only by AD cases (about 90%) and advanced LMCI participants (about 40%) but not by controls, this medication class largely coincides with diagnosis, leading to a highly significant correlation between medication status and diagnosis. As mentioned, we intentionally excluded diagnosis as covariate in regression analyses because the investigated clinical variables naturally also show high levels of correlation with diagnosis (and, thus, also with these medications).

For each metabolite, using a linear regression, we regressed metabolite level on medications. Medications were backward-selected via Bayesian information criteria to select an optimal combination of medications for preventing confounding while limiting model complexity. We refer to the residuals of each metabolite as the medication-adjusted metabolites. The residuals then carried forward to test associations with clinical and imaging outcomes.

### 2. ROS/MAP brain and serum metabolomics

Metabolomics was completed in 560 ROS/MAP participants. Brain and serum metabolomics was available for 93 participants. Human brain tissue and serum samples stored at -80°C (Forma 8600 series ultra-low temperature freezer, Thermo-Fisher Scientific, Nashville, NC) until sample preparation and analysis. Quantification of bile acid concentration was performed at the University of Hawaii cancer center. The bile acid-free matrix (BAFM) was used to prepare bile acid calibrators. Extracts of brain tissue along with bile acid reference standards were subjected to instrumental analysis. A Waters ACQUITY ultra performance LC system coupled with a Waters XEVO TQ-S mass spectrometer was used for all analyses. Chromatographic separations were performed with an ACQUITY BEH C18 column.
